# High-throughput creation and functional profiling of eukaryotic DNA sequence variant libraries using CRISPR/Cas9

**DOI:** 10.1101/195776

**Authors:** Xiaoge Guo, Alejandro Chavez, Angela Tung, Yingleong Chan, Ryan Cecchi, Santiago Lopez Garnier, Christian Kaas, Eric Kelsic, Max Schubert, James E. DiCarlo, James J. Collins, George M. Church

## Abstract

Construction of genetic variant libraries with phenotypic measurement is central to advancing today’s functional genomics, and remains a grand challenge. Here, we introduce a Cas9-based approach for generating pools of mutants with defined genetic alterations (deletions, substitutions and insertions), along with methods for tracking their fitness *en masse*. We demonstrate the utility of our approach in performing focused analysis of hundreds of mutants of a single protein and in investigating the biological function of an entire family of poorly characterized genetic elements. Our platform allows fundamental biology questions to be investigated in a quick, easy and affordable manner.

## INTRODUCTION

Libraries of cells with defined genetic alterations have proven transformative for connecting poorly understood genes to biological pathways and in uncovering novel roles for previously characterized genes. Despite these benefits, the high cost of development and requirement for extensive amounts of time and labor have meant that only a handful of such libraries have been developed. Furthermore, even in some of the most widely used collections, such as the yeast knockout library, a majority of the members contain undesired secondary mutations^1^ and the occurrence of false positives during screening due to the presence of selection markers within the genome affecting nearby genes^2^. As we continue to push the boundaries of genomics, there is a pressing need for scalable methods that enable the quick and cost-effective creation of multitudes of cells each with precise genetic alterations. These technologies would be further aided if the transition from library generation to library screening could be seamless, such that all of the generated variants could be screened within a single experiment *en masse*. In this work, we present a Cas9-based strategy for the simultaneous creation of hundreds of genetic variants within wild-type yeast cells that express the Cas9 protein along with a donor repair template. Using our platform, we generate two libraries targeting the DNA helicase SGS1: one containing a set of small tiling deletion mutants that span the length of the protein and another encoding a series of point mutations against highly conserved residues within the protein. We use these mutant libraries to characterize in detail the domain architecture and functional residues within SGS1 that are critical for its role in genome maintenance. Next, we create a genome-wide library targeting several hundred poorly characterized small open reading frames (smORFs, defined as ORF <100 amino acids in length) scattered throughout the yeast genome, and assess which are vital for growth under various environmental conditions in a single set of experiments.

## RESULTS

Our system is built upon CRISPR/Cas9 previously shown to be capable of enhancing genome editing in *Saccharomyces cerevisiae* via stimulation of homology-directed recombination (HDR) repair upon causing a double-stranded break at a given target locus^3^. Each isogenic mutant is generated by introducing a plasmid containing a targeting guide RNA (gRNA) paired with a corresponding donor template that contains the intended mutation we wish to introduce (hereon referred to as the guide+donor strategy) (**Figure 1A**). The advantages of our concatenated guide+donor design are threefold: a) It enables rapid cloning of all library members within a single set of reactions, b) it allows simultaneous delivery of both the guide and donor in one contiguous unit thus preventing uncoupling that may result in inefficient repair and unproductive repair outcomes, and c) it enables high-throughput molecular phenotyping using next generation (NGS) sequencing with each guide+donor containing a plasmid serving as unique barcodes for tracking edited cells. A similar concept of *in cis* delivery of guide+donor was recently demonstrated in bacteria^4^.

**Figure 1.**
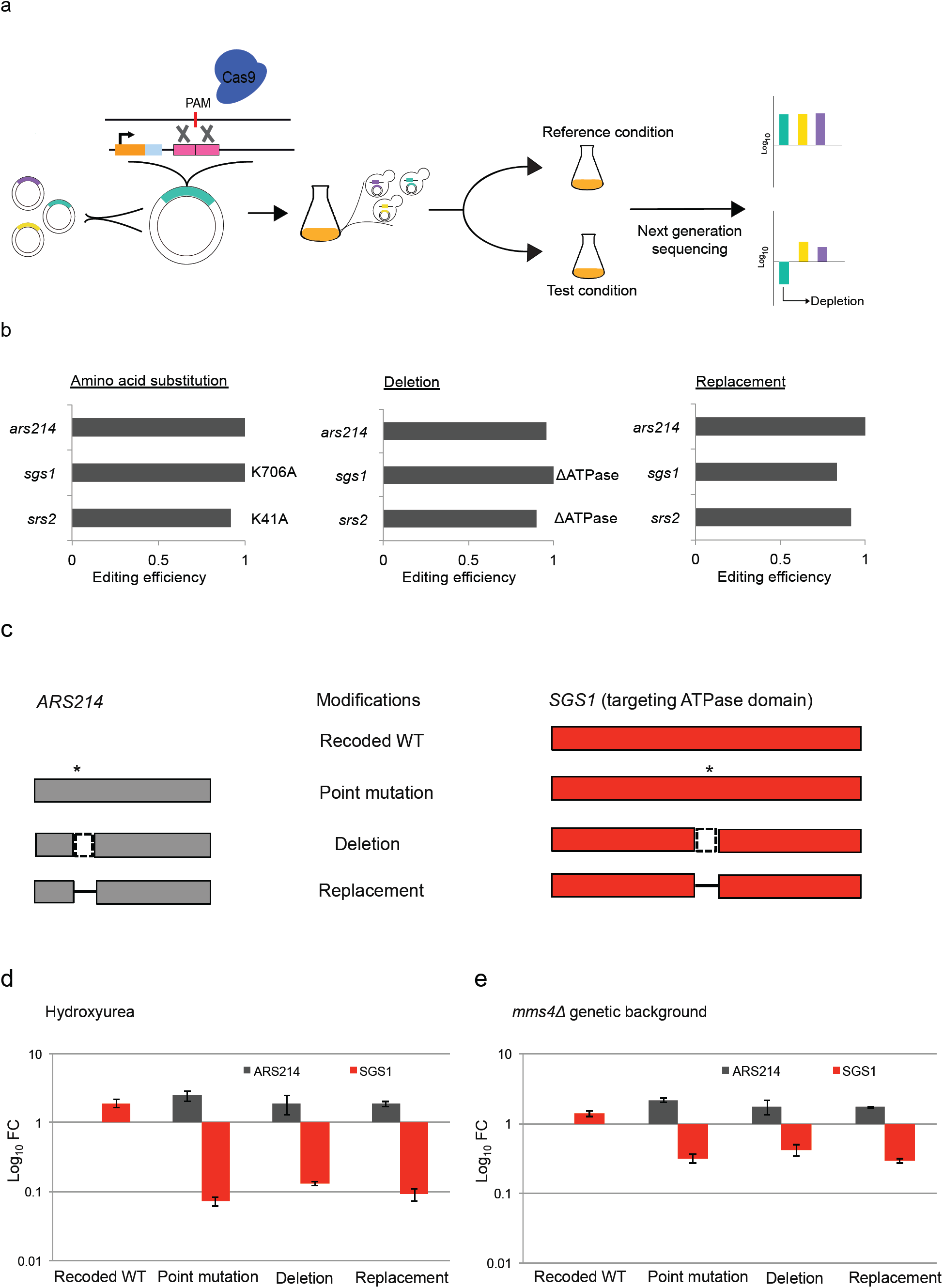
Guide+donor genome-editing platform to engineer and phenotypically characterize programmed mutations individually and in pools of mutants. (a) Schematic representation of guide+donor workflow. Briefly, guide+donor plasmids targeting different endogenous sites are indicated by different colors. Each guide+donor structure contains an *SNR52* promoter (orange), an N20 sequence (blue), a structural sgtail (not shown) and a donor template that contains desired mutations flanked by regions of homology (pink). Pools of guide+donor plasmids were simultaneously transformed into yeast. Libraries of yeast transformants were recovered and subjected to reference and test conditions. Genomic DNA from each sample was extracted, amplified, and subject to next generation sequencing (NGS) to detect depletion or enrichment. (b) Bar graphs showing genome editing efficiencies to generate various types of mutations (amino acid substitution, deletion, and sequence replacement) at three different endogenous sites (*ARS214, SGS1, and SRS2*). Catalytic amino acid substitutions and ATPase domain deletions for *SGS1* and *SRS2* are indicated. (c) Schematic representation of programmed edits in *ARS214* and *SGS1* generated via guide+donor in pool for phenotypic characterization in (d) and (e). Asterisk, dotted box, and solid dash denote substitution, deletion, and replacement of an amino acid stretch with a linker sequence, respectively. Variants of *ARS214* and *SGS1* are indicated by colors gray and red, respectively. Figures not drawn to scale. (d) Bar graph showing fold change of guide+donor library members targeting *ARS214* and *SGS1* in response to HU treatment. Types of genetic modifications are indicated on the x-axis. Depletion is represented in log scale on the y-axis. Data are shown as mean ± s.e.m. (n = 2 independent yeast transformations) (e) Bar graph showing depletion of *ARS214* and *SGS1* guide+donor library members in *mms4δ* genetic background. Genetic modifications and their corresponding abundance in log scale are indicated on x- and y-axes, respectively.

As a test of our guide+donor system, we integrated a copy of the Cas9 gene into the neutral *HO* locus and then transformed in a guide+donor containing plasmid designed to disable the *ADE2* locus. However, upon selecting for cells with the guide+donor, we found that the number of colonies with the desired genetic alteration were low (<5%), consistent with earlier attempts at *in cis* guide+donor delivery in yeast^5^. We sought to increase the percentage of correctly edited cells in order to enable efficient genome-scale measurements via NGS.

Previous work has shown that linearized donors can boost conversion efficiency by 10-1000 fold compared to closed circular donors^6-12^. To test if linearization of our guide+donor plasmid would increase the efficiency of our system, we introduced our guide+donor system as two linear pieces of DNA. The larger DNA fragment contained the guide+donor portion of the plasmid, along with an internal portion of the selection marker removed. The smaller DNA fragment consisted of the missing portion of the selection marker along with ∼150 bp of flanking homology such that HDR was required to reconstitute the full circular plasmid (**Supplementary Figure 1A**). When this modified approach was employed, we observed a marked increase (>90%) in the number of transformants and colonies with the desired repair event (**Supplementary Figure 1B**).

To begin characterizing the limitations of our system, we tested a series of vectors designed to introduce either targeted point mutations, short deletions or sequence replacements within the *ADE2* locus. For programmed point mutations, we obtained an efficiency of genome modification close to 100% for changes that occurred proximal to the Cas9 generated cut site (**Supplementary Figure 2A**). In contrast, when the desired mutation was placed further away from the Cas9 cut site, we noted a decrease in the efficiency of our system, with mutations 12- 15 bp away showing rates of editing of ∼40%. Next, we tested our ability to delete a set of contiguous bases; using our approach, we were able to remove up to 61 bp at a time with high efficiency (>90% of colonies with the desired change), but larger deletions (≥121bp) showed a sharp decline in efficiency (<25%) (**Supplementary Figure 2B**). Finally, we examined our ability to perform blocks of contiguous sequence replacement by deleting 61bp of an endogenous site and replacing it with up to 15bp of desired sequence and observed levels of repair similar to performing the deletion alone (>90% of colonies with the programmed change) (**Supplementary Figure 2C**).

Having gained insight into the limitation of our guide+donor strategy, we next sought to determine the generality of our method by targeting three additional loci (*SGS1, SRS2* and *ARS214*) with a series of point mutations, deletions and sequence replacements. Similar to our *ADE2* results, we obtained a high efficiency of genome modification (90-100%), across all targets and mutation types (**Figure 1B**).

The strong correlation between the presence of a particular guide+donor plasmid within a cell and the cell having the desired genetic alteration allowed us to test if, by sequencing the abundance of different guide+donor pairs within a mixed pool, we could infer the fitness effects of their modifications. To test this possibility, we built a small library containing a mixture of guide+donor plasmids designed to modify either the non-essential *ARS214* locus or the DNA damage repair helicase *SGS1* (**Figure 1C**). The cells that arose from the pooled transformation were then subjected to growth in media with or without the genotoxic agent hydroxyurea (HU) and the abundance of the various guide+donor plasmids within the population was determined by high-throughput sequencing. As hypothesized, we observed a marked depletion of guide+donor pairs that encoded modifications that disrupted the ATPase domain of Sgs1, which is known to play a critical role in its function (**Figure 1D**). In a second mock library, we had guide+donor pairs designed to mutate the C-terminus of *SGS1*, which has been shown to lead to a milder loss of function phenotype. In line with our expectations, the C-terminal mutants showed less depletion (**Supplementary Figure 3)**. As a control, our small library also contained guide+donor pairs that introduced synonymous changes within the ATPase domain or C-terminus of *SGS1*. As expected, these pairs did not show depletion, suggesting that the effects observed were due to cells receiving the intended modifications and not due to non-specific disruption of the *SGS1* locus by Cas9. Furthermore, when each of the generated strains was tested individually, the results correlated well with our pooled analysis, lending additional support for the validity of our method (**Supplementary Figure 4**). Along with testing the effects of environmental perturbation on our mutant library, we also examined if our system could be used to observe gene-gene interactions by transforming our small library into cells defective in the structural endonuclease Mms4. Upon transformation of the same library into an *mms4δ* genetic background, we observed ∼5-fold depletion across all *sgs1* mutants, consistent with known synthetic sickness between *SGS1 and MMS4* (**Figure 1E**).

Having established the ability of our system to perform one-pot genome engineering followed by *en masse* characterization, we sought to scale up our approach and apply it towards the systematic characterization of a single protein, Sgs1, or the genome-wide analysis of a family of hundreds of poorly characterized small open reading frames.

To date, many gene-based studies have determined the role of a given protein by examining cells in which the protein is entirely absent. While informative, these studies are often poor surrogates for modeling the targeted loss of function mutations that are observed within patient populations. Furthermore, due to their gross nature, these studies also fail to provide insights into which of the many biological activities carried out by a single protein are central to the observed phenotype. To test if our system might be used to gain a more thorough understanding of protein activity, we generated two libraries against *SGS1*, encoding the yeast homologue of the human DNA helicase BLM with known roles in mitotic stability, cancer, and aging^13^. To map the domains within the Sgs1 protein that are critical in providing cellular resistance to the genotoxic stressor HU, we designed a set of guide+donor constructs that generate 20 amino acid deletions with 5 amino acid sliding windows across the majority of the *SGS1* gene (see Materials and Methods for details). Among the regions showing strongest depletion within edited cells were guide+donors deleting amino acid stretches 1-85, 686-1090 and 1116-1225 within Sgs1, which correspond to the Sgs1-Top3-binding domain, Sgs1-helicase, and RQC domains, respectively (2-tailed t-test, P<0.0001) (**Figure 2, Supplementary File 1**). These results are consistent with the known mechanism by which Sgs1 functions through the recruitment of accessory proteins (through N-terminal residues) ^14-22^ and by resolution of DNA structural intermediates via its helicase and RecQ domains^23,24^.

**Figure 2.**
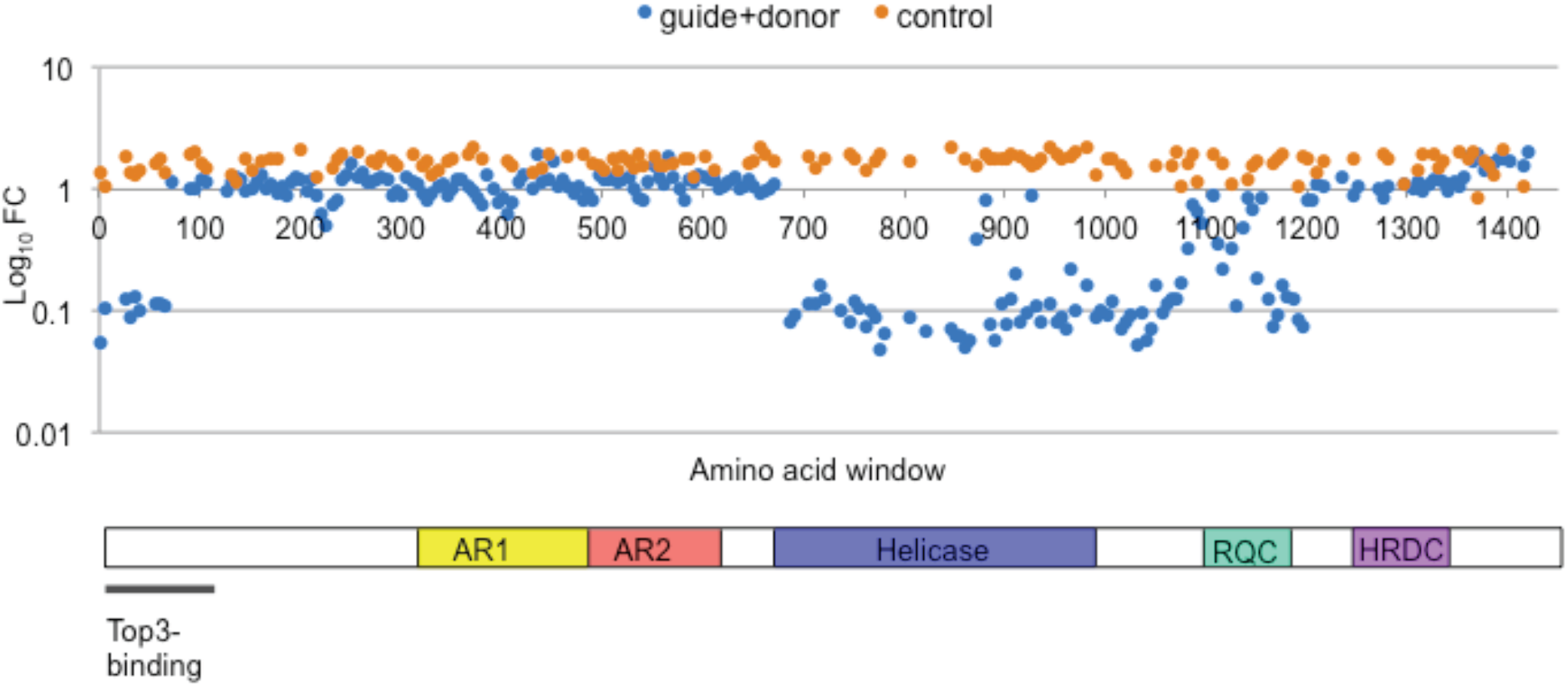
Guide+donor library of *sgs1* mutants in response to HU. (a) Scatterplot showing fold change in abundance of guide+donor members programmed to generate *sgs1* tiling deletion mutants across the entire *SGS1* gene in response to HU. Guides paired with corresponding donor sequences to generate programmed deletions are indicated in blue. Guides paired with sequence that lack homology regions to qualify as donors, defined as junk donors, are indicated in orange. X-axis denotes the amino acid window along the protein. Level of depletion represented in log scale is indicated on the y-axis. Schematic representation of relevant domains and motifs in Sgs1 is shown. Figures not drawn to scale.

Next, we used our method to create a series of precise point mutations within Sgs1. Towards this goal, we selected a set of evolutionarily conserved amino acid residues within the Sgs1 helicase domain and changed them to all other possible amino acids using our guide+donor strategy. This library was then exposed to growth in HU to assay for mutant drug sensitivity. Despite targeting highly conserved residues within Sgs1, all but one tolerated alanine substitution without causing an obvious loss in resistance to our highest concentration of HU at 40 mM (**Figure 3, Supplementary File 2**). In the case where activity was lost, it was when alanine was used to replace the essential helicase catalytic residue K706. Examining our data as a whole, we observed expected trends of amino acid substitutions of similar charge and size being well tolerated while those with opposite properties being more detrimental to Sgs1 function.

**Figure 3.**
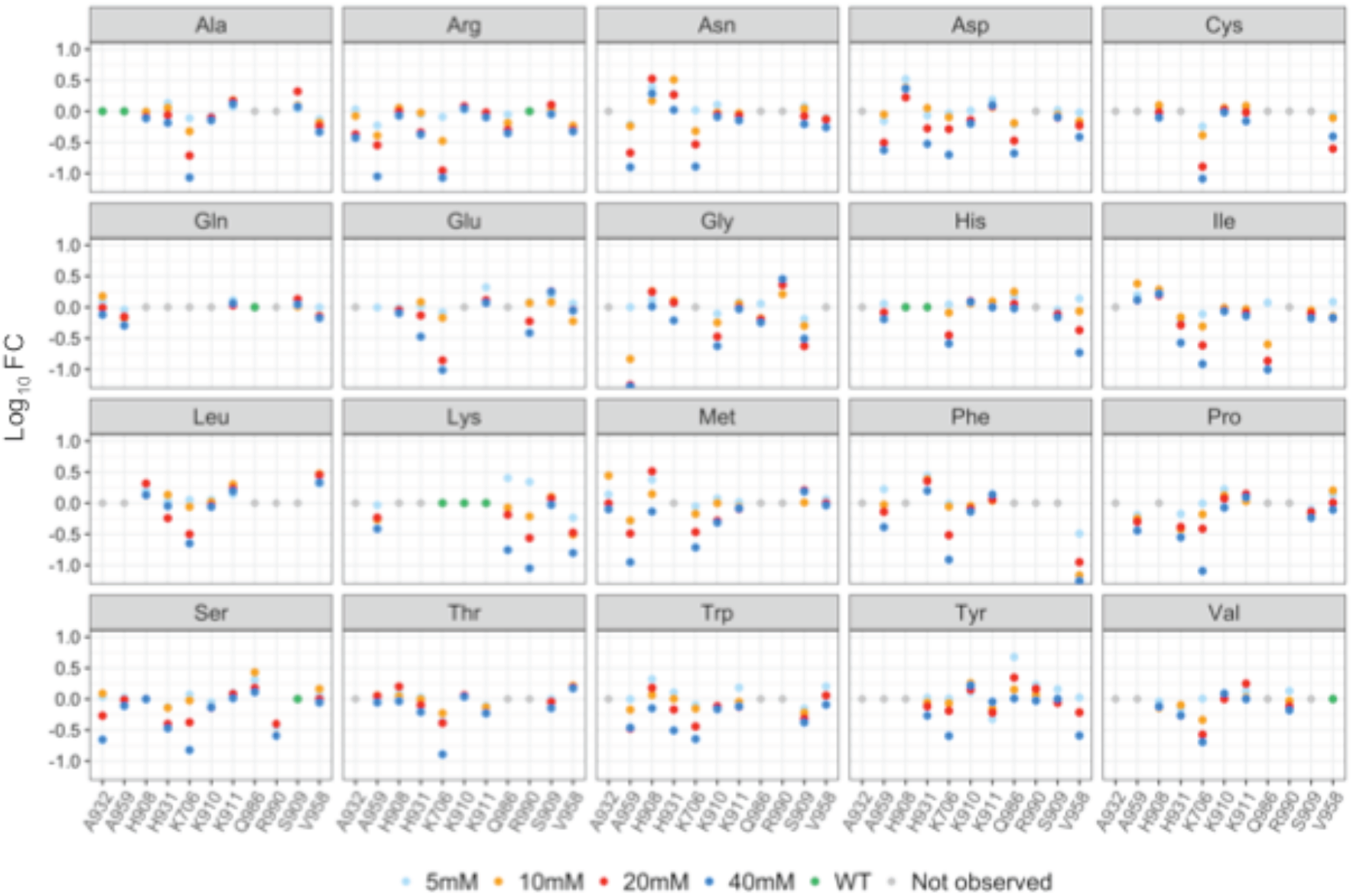
Guide+donor library of amino acid substitutions of selected conserved residues in *SGS1* in response to various concentrations of HU. Scatterplots showing fold change in abundance of guide+donor members programmed to generate precise point mutations within Sgs1 in response to HU. Various conditions of HU are represented by different colors in the legend. Selected conserved residues are displayed on the x-axis. Y-axis denotes fold depletion in log_10_ scale. Each subplot shows the corresponding amino acid to which each conserved residue was replaced.

Given the success of our platform in building libraries of mutants against a single gene, we sought to determine its capacity to perform targeted editing across the entire yeast genome. As a proof-of-concept, we built a guide+donor library designed to cause small deletions around the initiating ATG for a set of randomly chosen canonical ORFs (including both essential and non-essential genes) along with several hundred poorly characterized smORFs. Unlike canonical ORFs, smORFs remain largely ignored and are often missing in modern genome annotations due to their size, low conservation scores, and lack of similarity to known proteins and protein domains.

Using our genome-scale deletion library, we first analyzed which of the targeted members were essential for viability. We observed strong depletion (∼3- 10 fold) for all targeted essential ORFs (2-tailed t-test, P<0.0001) compared to <2-fold depletion for nearly all nonessential ORFs (2-tailed t-test, P=0.18), thus highlighting the specificity and sensitivity of our method (**Figure 4A, Supplementary File 3**). Out of the hundreds of smORFs that were examined, 20 smORFs showed similar levels of depletion as our essential controls (2-tailed Z test, P<0.0018) with previous results^25^. Furthermore, although a number of the smORF library members are located in close proximity upstream to a nearby essential ORFs (in some cases within 132 bp), our screen did not identify any of them as essential, providing additional evidence for the specificity of our targeting method. To further demonstrate the ability of our guide+donor strategy to characterize a large number of proteins in parallel, we subjected our smORF mutant library to a series of environmental stressors including growth: at 37°C (**Figure 4B**), in the presence of HU (**Figure 4C**), or with the antifungal drug fluconazole (**Figure 4D**). For each of our screens, we identified nearly all of the previously known smORFs with tolerance towards each of the tested conditions, along with uncovering novel roles for a large number of additional smORFs^26^.

**Figure 4.**
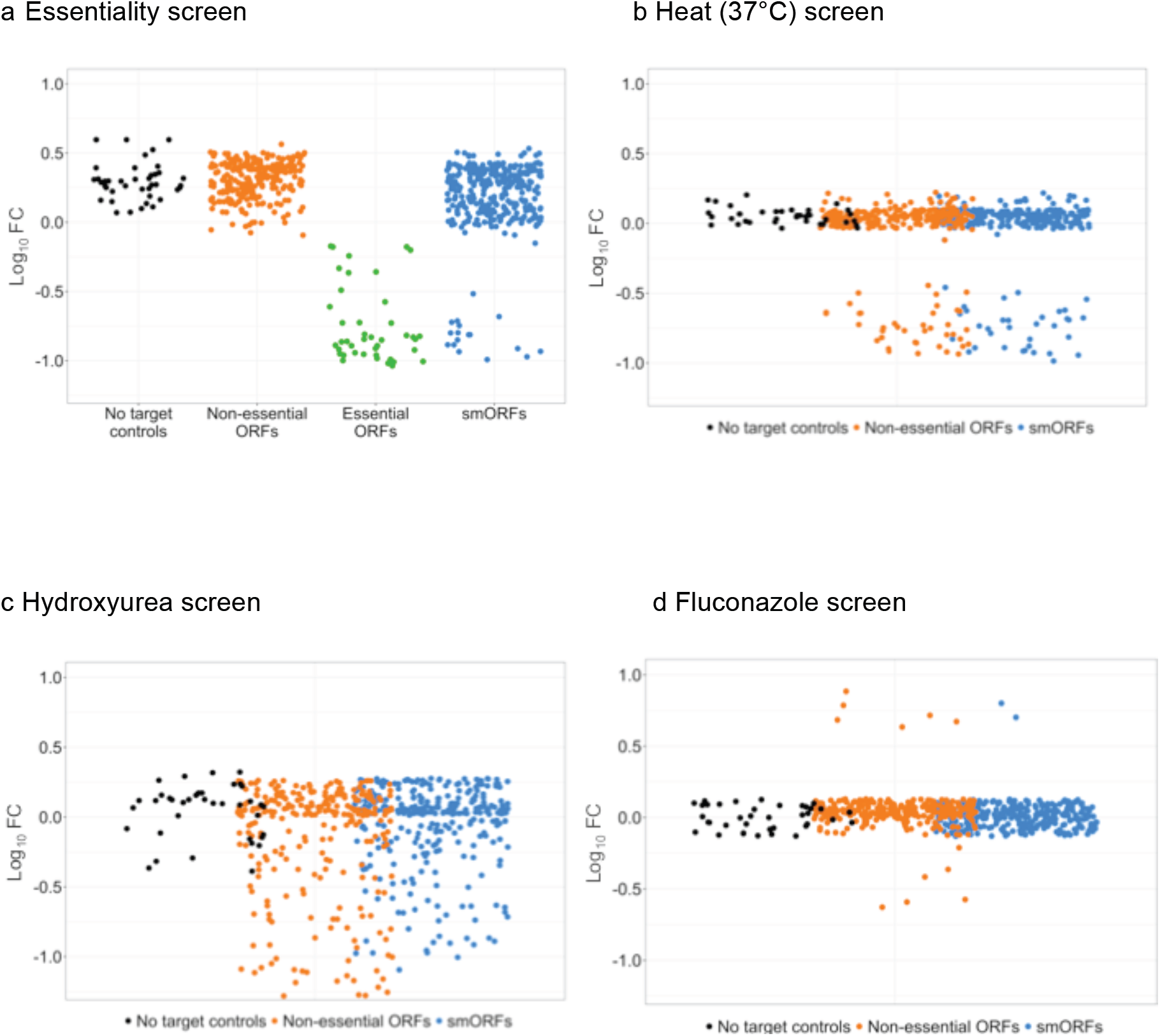
smORF mutant library subject to different phenotypic screens. (a) Scatterplot showing fold change in the abundance of guide+donor reads based on essentiality of the corresponding mutant. Guide+donors categorized into different groups based on gene targets are indicated on the x-axis. Degree of depletion, in log_10_ scale, is shown on the y-axis. (b) Scatterplot showing sensitivity of only non-essential library mutants exposed to 37°C heat. Gray, orange and blue dots represent mutant cells containing guide+donor plasmids that are no-target controls, targeting ORFs, or targeting smORFs, respectively as shown on the x-axis. Y-axis denotes fold depletion in log_10_ scale. (c) Scatterplot showing sensitivity of only non-essential mutant members to HU. Same axes labels as in (b). (d) Scatterplot displaying log_10_ fold change of non-essential targets in response to anti-fungal fluconazole. Same axis labels as in (b)

Of the 315 smORFs that were screened, 66 were found to play a role in cellular fitness under the tested conditions. This is in stark contrast to conventional ORFs for which nearly twice as many were found to be involved in growth under the same environmental conditions (112 of 308 tested ORFs) (Chi-square test, P<0.0001). We next sought to determine if there were any properties shared by the smORFs, which showed biological activity, as the insights gained might enable later groups to better annotate genomes for these biologically significant elements. Although smORFs show a range of sizes across the yeast genome (smallest smORF hit was 29 aa), we found that as a whole, longer smORFs along with those showing elevated levels of RNA expression exhibited a trend of slightly more likely to come up as hits in our screen (**Supplementary Table 1**). Of note, ORFs showed no such correlation with regard to length, but maintained a similar trend with respect to expression (**Supplementary Table 2**). We next used computationally derived predictions of protein secondary structure to test if any variation in the prevalence of structural elements (e.g. alpha helices and beta-sheets) existed within smORF hits as compared to non-hits. No difference in the prevalence of secondary structural elements were seen between smORF hits vs non-hits, yet we did observe an increased propensity for beta-sheets and a decrease in unstructured loops when smORFs as a whole were compared to the set of ORFs that were also examined in our screens (**Supplementary Table 3**). Finally, as evolutionary conservation is a strong predictor of biological importance, we determined the rate of conservation across multiple species between smORF hits within our screens and those that showed no activity (**Supplementary Table 4**). A large difference in the rate of gene conservation was found with 29 of the 66 smORF hits being conserved in humans as compared to only 44 of the 250 nonfunctional smORFs (Chi-square test, P<0.0001).

## DISCUSSION

Here, we present a high-throughput method for the rapid generation and phenotypic characterization of hundreds of mutants and illustrate its potential in domain/residue mapping and functional interrogation of nearly any user-defined genomic target. To understand the strengths and limitations of the guide+donor method, we empirically determined the design parameters required to efficiently generate programmed mutations, namely deletions, amino acid substitutions, and sequence replacements. This enables the creation of specific user-defined loss-of-function, gain-of-function, and altered regulation mutants *en masse*. More importantly, by editing the locus within its native context without the need for exogenous marker genes, we avoid artifacts from using surrogate reporter systems and false positive and negative results due to selection marker-driven positional effects^27^. Furthermore, the fact that our system is highly efficient allows users to simply read the guide+donor sequence present within the plasmid delivered to each cell and, by doing so, identify the cell’s genotype. Ultimately, this feature enables the fitness of hundreds of mutants and potentially thousands to be tracked by simply sequencing the abundance of each guide+donor sequence within a population.

We validated our platform using the well-characterized DNA helicase *SGS1*. Upon verifying its accuracy, we applied our technology to examining the roles of hundreds of poorly characterized smORFs. To our knowledge, our *SGS1* tiling deletion data represents the first comprehensive domain bashing of a eukaryotic protein, thus showcasing the ability of our technology to rapidly hone in on the critical domains required for protein function. Kastenmayer *et al*. (2006) used conventional techniques to make specific gene deletions of 140 smORF mutants^25^. In contrast, we demonstrated the ease of our guide+donor method in rapidly covering over ∼77% of the 299 putative smORFs within the yeast genome, including many that had previously been neglected^25^. Importantly, our ability to capture most known hits identified from previous work established the specificity and sensitivity of our platform in systematic interrogation of gene function. Given the degree of conservation between yeast and human genomes and the conservation between several smORFs and higher eukaryotes^25^, it will be interesting to see if the smORFs identified in our work with roles in stress tolerance have similar functions in humans.

In our studies, we have employed the commonly used *Streptococcus pyogenes* Cas9 (SpCas9) to catalyze our guide+donor system. Because SpCas9 requires the availability of an NGG protospacer adjacent motif (PAM) at the target region, our system is currently bounded by this limitation. However, SpCas9 variants recognizing alternative PAMs such as NAG or NGA and Cas9 proteins from other species which use alternative PAMs are also available and, in principle, should be readily employed within our system to greatly broaden the range of sequences that can be modified by our platform^28^.

*S. cerevisiae* remains a major lab workhorse and the model system through which much of our understanding of eukaryotic cell biology is derived. While we have focused on the usage of our technology for the high-throughput characterization of coding elements, we envision a broad range of additional applications such as: directed evolution, metabolic engineering, and in exploring the function of non-coding elements (both RNAs and regulatory sequences). Moreover, given that most clinically relevant mutations are point mutations and not gene knockouts, along with the high degree of gene conservation between yeast and humans, our guide+donor editing platform provides an easy way to engineer and test the effects of hundreds of currently uncharacterized single nucleotide polymorphisms that exist within human populations via their nearest yeast ortholog. We envision that this kind of study may enable the first practical genome-wide assessment of the effects of previously uncharacterized missense mutations within the human population and, by doing so, provide helpful insights into causal mutations while allowing an improved understanding of underlying disease mechanisms.

## ACKNOWLEDGEMENTS

G.M.C was supported by NIH grants RM1 HG008525 and P50 HG005550. A.C. was funded by the National Cancer Institute grant no. 5T32CA009216-34. J.J.C was funded by the Defense Threat Reduction Agency grant HDTRA1-14-1-0006, the Paul G. Allen Frontiers Group. X.G. and A.C. conceived of the idea, led the study and designed all experiments. A.C. and R.C. with input from J.E.D. demonstrated the initial feasibility of the guide+donor approach. X.G. performed the majority of experiments, including the oligo library design, library construction and analysis with significant technical contribution from A.T. R.Y.C. provided expertise in statistical analysis. S.L.G. assisted with oligo library design and NGS data analysis. C.K. generated the RNA-seq data for BY4741 yeast strain and provided the FPKM values. E.K. provided insight with regard to library construction methods and analysis. M.S. provided technical expertise with regard to methods to increase guide+donor efficiency. J.J.C and G.M.C. oversaw the study. X.G. and A.C. wrote the manuscript with input from all authors.

